# OmicsNet: Integration of Multi-Omics Data using Path Analysis in Multilayer Networks

**DOI:** 10.1101/238766

**Authors:** Murodzhon Akhmedov, Alberto Arribas, Roberto Montemanni, Francesco Bertoni, Ivo Kwee

## Abstract

Integrative analysis of heterogeneous omics data is essential to obtain a comprehensive overview of otherwise fragmented information and to better understand dysregulated biological pathways leading to a specific condition. One of the major challenges in systems biology is to develop computational methods for proper integration of multi-omics datasets. We propose OmicsNet that uses a multilayer network for the integration and analysis of multi-omics data of heterogeneous types. Each layer of the multilayer network represents a certain data type: input layers correspond to genotype features and nodes in the output layer correspond to phenotypes, while intermediate layers may represent genesets or biological concepts to facilitate functional interpretation of the data. OmicsNet then calculates the highest coefficient paths in multilayer network from each genomic feature to the phenotype by computing an integrated score along the paths. These paths may indicate the most plausible signalling cascade caused by perturbed genotype features leading to a particular phenotype response. With example applications, we illustrate the potential power of OmicsNet in the functional analysis, biomarker discovery and drug response prediction in personalized medicine using multi-omics data.

## BACKGROUND

### Integration of omics data

The development of effective therapeutic approaches may be facilitated by a system-level understanding of the molecules altered in a disease, as well as their complex interactions among themselves. To understand the pathogenesis of a disease, it is crucial to analyze underlying mechanisms causing the changes in tumor versus normal patients. Systems biology is emerging field in this aspect, which systematically measures the molecular perturbations within the cell and targets to elucidate the features and dysregulated biological pathways resulting the disease. Various experiments may be used to measure alterations in DNA, RNA, protein, and metabolites levels between disease and control systems. With the recent developments and reduced cost in high-throughput technologies (Mias and Snyder 2013), a vast amount of data including the copy number, mutation, methylation, gene expression, proteomic, and metabolomic data at multiple layers of cellular systems can be profiled. Even though these omics data is obtained by measuring the perturbations in the same cell, the molecular information is fragmented over layers in each data type and overlap is considerably small, because they narrate the complementary parts of the same biological process. In order to have a complete picture there is a need for methods that properly *integrate* and interpret multi-omics datasets. Rather than analyzing data types separately, analyzing *multi-omics* and phenotypic sources of information jointly could enhance our understanding of complex circuitry models of interaction in new ways. The current bioinformatics challenge is to tackle the problem of fragmentation of knowledge by integrating several source of data into one entity.

It is not trivial to consolidate different types of genomic data due to different scales and noise in each dataset, the high number of features compared to sample sizes, complementary nature of the datasets, and the complexity of biological processes (B. Wang et al. 2014). A wide range of mathematical methods have been proposed for the integration of multi-omics data, for a full review of methods see Bersanelli et al. (2016). Ritchie et al. (2015) broadly classifies integration methods into multistaged and multi-dimensional approaches, and further subclassifies the latter into concatenation-based, transformation-based and model-based integration methods.

### Multilayer networks for omics data integration

Recently, multilayer and multiplex networks have been proposed to analyze and integrate high-throughput data (Bersanelli et al. 2016). A multilayer network is composed of multiple layers, which can be interpreted as a ‘layers of networks’. Each layer corresponds to one data type. Vertices in the layers corresponds to gene features with intralayer connections to other features in the same layer, and interlayer linkage to other layers. Integration using multi-layer networks can be characterized as a combination of multi-staged and model-based integration methods.

Most of the existing approaches in biological context use the multilayer networks mainly for clustering to stratify patients through integrative analysis of omics data. Similarity Network Fusion (SNF) is a network fusion tool, which constructs an integrated patient-similarity network based on each data type separately, and effectively fuses multiple networks into a single network that demonstrates the full spectrum of underlying data using the complementary nature of different sources (B. Wang et al. 2014). Tested on five different cancer types, SNF outperforms the single data type analysis, demonstrating the power of integrative analysis. A similar fusion approach was developed further fusing transcriptomic and fluxomic layers using a multiplex network, capturing similarities across two omic levels (Angione et al. 2016). Modularization of multilayer networks has been used to find consensus signature modules across the data types that are functionally associated with targets in aging (Dimitrakopoulou, Vrahatis, and Bezerianos 2015). iCluster is a joint latent variable model developed for flexible modeling of the relations between different data types and clustering of patients (Shen, Olshen, and Ladanyi 2009). A recent approach adapted the generalized modularity technique for optimizing the partitioning in patient-similarity networks (H. Wang et al. 2016). The main application of the method was characterization and identification of cancer subtypes. Gao et al. (2015) presented a method to reverse engineer integrative gene networks that works by integrating different quantitative and qualitative data sets by reconstructing a multiplex network and solving small quadratic programs based on local neighborhood of nodes. A computational method on multiplex networks based on Levy random walks has also been presented, but not supported by application on biologic data Guo et al. (2016). Also other methods have been published, but none of them includes the use of the path analysis or shortest path algorithm (Castellani et al. 2014; J. Kim, Woo, and Nam 2016). Finally, other studies use multiplex networks to understand the molecular mechanism behind the biological processes by integrating heterogeneous omics data (Moni and Liò 2015; Bersanelli et al. 2016; Yugi et al. 2016) or used multilayer and multiplex networks to model complex interactions in social networks, physics, and computer science (Kivelä et al. 2014; Menichetti et al. 2014; De Domenico et al. 2016).

### Proposed multilayer framework for multi-omics data integration and analysis

We present OmicsNet multilayer framework for integrative analysis of heterogeneous omics data. Our data integration method is based on the idea to model the multi-omics data as a multilayer network and then subsequently perform path analysis in this network. One layer can act as enhancer or inhibitor of the other. For instance, high methylation can inhibit the gene expression and reciprocally high copy number gain may enhance it. To the best of our knowledge, three features of our approach are novel.

- First, we use multilayer networks where the specific configuration of the layers is based on plausible flow of the biological processes as shown in Fig. 1. For example, the copy number and methylation layers are in front of gene expression since they regulate the expression. The information flows from the input layers to the output layers as shown by directed edges.
- Second, we add functional layers such as geneset and so-called ‘biological concept’ layers to better elucidate through which higher order functional mechanisms the genotype perturbation cause the phenotype. Note, the data in these layers are computationally derived from the measured data.
- Then, after assigning weights to the edges and prizes to vertices, we compute the maximum coefficient path (or set of paths) in the multilayer network that have the highest feature correlation to the phenotype. The set of nodes and edges on these paths facilitate to understand the signaling cascades that are being altered from perturbed features to the cellular response.

**Figure 1:**
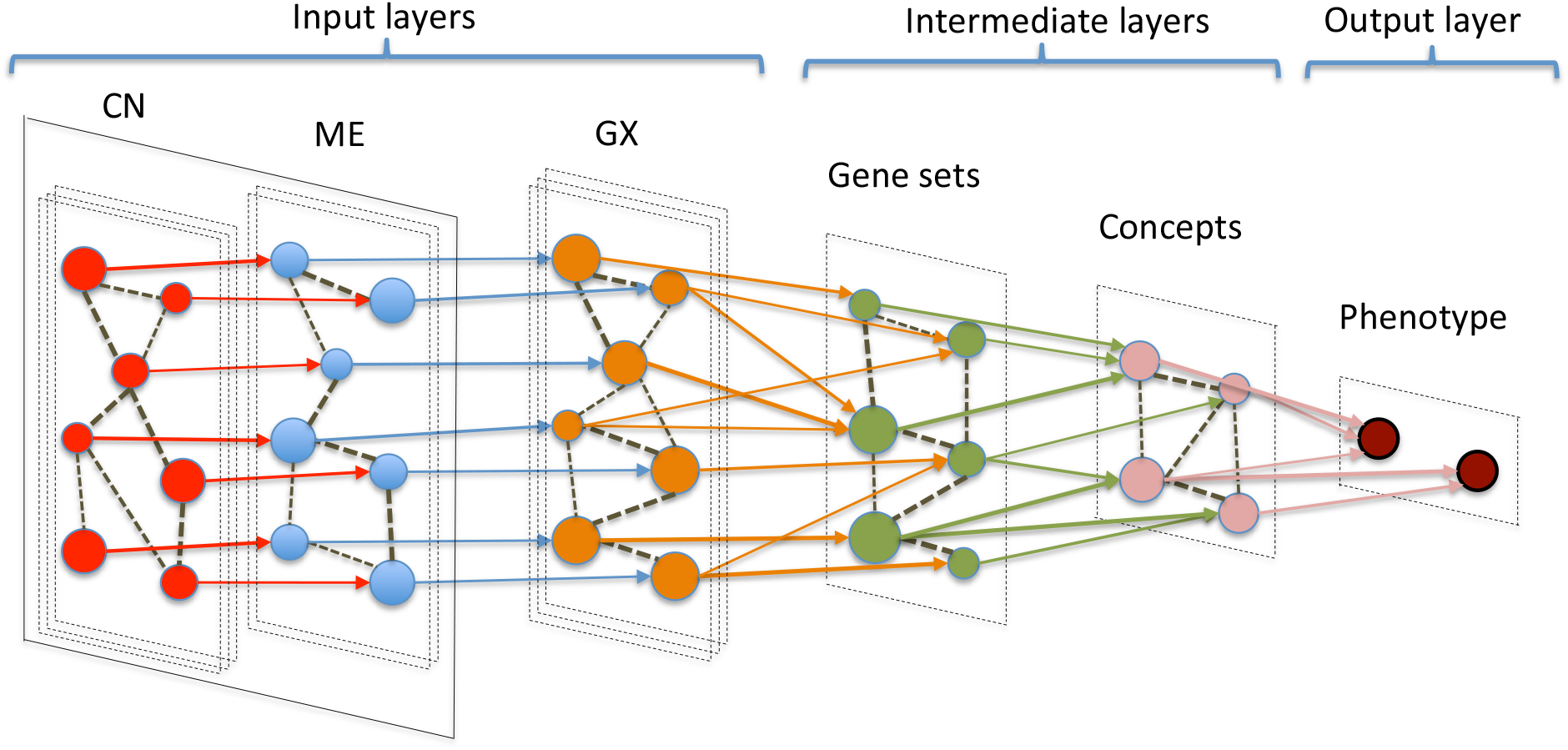
The proposed multilayer network for integration of heterogeneous omics data composed of input, intermediate and output layers. The information propagates from the left (input layers) to the right (output layers). Abbreviations used for labeling the layers are CN = copy number, ME = methylation, and GX = gene expression. The intra-layer edges are represented by dashed lines and inter-layer edges connecting the two layer are shown in solid lines.

Existing multilayer methods as well as our proposed method are summarized in Table 1 and compared against each other in terms of network structure, employed algorithm and the application area.

**Table 1:** Summary of omics integration methods using multilayer network and their main algorithm/application.

| reference | network | algorithm | application |
| --- | --- | --- | --- |
| **(our method)** | **multilayer** | **max-coef path\*** | **integrative analysis** |
| Angione et al. (2016) | multiplex | SNF | **integrative analysis** |
| B. Wang et al. (2014) | multiplex | SNF | clustering |
| H. Wang et al. (2016) | multiplex | gen modularity | **integrative analysis** |
| Gao et al. (2015) | multiplex | quadratic prog | network inference |
| Bersanelli et al. (2016) | *various* | *various* | **integrative analysis** |
| Guo et al. (2016) | multiplex | random walk | *not specified* |
| Moni and Liò (2015) | bipartite | **shortest path** | clustering |
| Yugi et al. (2016) | **multilayer** | *not specified* | **integrative analysis** |
| Castellani et al. (2014) | **multilayer** | *not specified* | **integrative analysis** |
| J. Kim, Woo, and Nam (2016) | **multilayer** | *not specified* | **integrative analysis** |

\*maximum coefficient path using the shortest path algorithm

### Path analysis in multilayer networks

The path analysis in multiplex networks, specifically the shortest path, has been studied in Ghariblou et al. (2017) to better define the concept of betweenness centrality and applied to find the most influential nodes in the multiplex setting. To our knowledge, no one has proposed applying the highest coefficient paths analysis in multilayer networks for genomics data integration and its analysis. Our network-based framework infers co-regulated genes and pathways from heterogeneous omics data by computing the set of maximum coefficient paths in weighted multilayer networks.

## METHOD OVERVIEW

*Note: Mathematical details and implementation notes are separately described in the Supplementary Methods of this paper*.

### Defining the multilayer network

OmicsNet implements a multilayer network with input, intermediate and output layers, where every layer represents a certain data type and nodes may represent a gene, geneset, or phenotype based on which layer it is located (**Fig. 1**). Layers can have both intralayers connections (connecting nodes within the layer) and interlayers connections (connecting to different layers). Input layers generally correspond to genotype features (measured omics types). Intermediate layers represent genesets, pathways and higher level concepts for biological interpretation of the data. The final output layer (or ‘phenotype layer’) corresponds to sample phenotypic information such as histology or cell type, or clinical information in the case of a patient, such as cancer subtype, drug response, or survival. The signalling information in multilayer network framework propagates from the input layers to the output layers as shown by directed edges in **Fig. 1**.

The number of layers is not limited and depends on the number of different data types and functional layers in the model. The specific configuration of the layers is based on plausible flow of the biological processes. For instance, copy number and methylation layers are modelled before the gene expression layer since they are assumed to regulate the expression.

For a particular layer, one may disallow intralayer connections to model *independency* between features. One may restrict the number of intralayer connections to improve computational and memory efficiency. Between the omics layers, interlayer connections may be restricted to nodes that belong to the same gene/protein to model CIS-regulation between data types. Note that the correlation values on the edges are independent of the selected phenotype but the vertex prizes not.

Intermediate layers generally correspond to other measured data types, or can be computationally derived layers from measured data types such as the gene set or biological concept layers. The biological concepts are composed of ‘set of gene sets’ and they provide higher-level functional information. The concepts decrease the number of gene sets to be analyzed and simplify the data interpretation. Geneset and concept layers are computed from the primary omics data types using summary methods such as GSVA, ssGSEA, or any other method that calculates single-sample gene set statistics (Tarca, Bhatti, and Romero 2013). These layers help to better elucidate through which functional mechanisms the genotype perturbation cause the phenotype.

The phenotype layer may contain multiple phenotype features, thus the multilayer model can be used for the joint analysis of multiple phenotypes. Intralayer connections in the phenotype layer may model multi-collinearity in the phenotype information.

Additional omics data types can be easily incorporated by adding additional layers. For instance, the proteomic, phosphoproteomic, and metabolomic data can be added between the gene expression and gene set layers since they harbor post-transcriptional and post-translational biological information.

### Scoring the multilayer network

Each intralayer and interlayer edge linking two features in OmicsNet multilayer framework is associated with a weight calculated based on the correlation between those features. We generally consider only positively correlated edge weights since it is easier to interpret the relationship between features and consequent phenotype. In certain cases, one may consider negative correlation value in the computations while modeling negatively correlated biological information flow. Otherwise, the absolute correlation value may be used as edge weights when the sign of the biological information flow is deemed to be irrelevant.

Each variable (vertex) is labeled with a ‘prize’ that quantifies some perturbation of that variable with respect to the phenotype. It can be computed by the correlation between that variable and phenotype. For instance, the prize of a vertex can be a copy number difference, expression fold change, number of mutations, or any parameter that quantifies some association with the phenotype. We generally take the absolute value of the prizes, or perform the path analysis for positive and negative node prizes separately.

### Computing the maximum coefficient paths

After assigning proper weights to the edges and prizes to vertices, OmicsNet computes the maximum coefficient path (**Supplementary Methods Eq. (2)**), or a set of highest coefficient paths (**Supplementary Methods Eq. (3)**), in the multilayer network. A simple illustration of the path coefficient calculation is shown in **Fig. 2**. Note that the maximum coefficient path between genotype and phenotype is constrained by the structure of the multilayer network, and generally must traverse all omics layers and higher order functional layers. The set of nodes and edges on these paths facilitate to understand the signaling cascades that are being altered from perturbed features to the cellular response. Computing the maximum coefficient path is NP-hard problem that requires exponential time for arbitrary graphs. However, we mathematically convert it into the shortest path (SP) problem (**Supplementary Methods Eq. (5)**) for efficient computation of optimal solutions. The shortest paths through the multilayer network can be efficiently computed by Dijkstra’s algorithm solving for single-source shortest path. The SP solution traverses each layer of the multilayer network at least once and in layers that have intra-layer connections, it is possible that the SP contains multiple traversals *within* a single layer. For each SP solution, the method calculates an associated *path coefficient*. In its basic form, this score is computed as the product of correlation values (**Supplementary Methods Eq. (1)**) and quantifies the joint correlation of all variables along the SP path from genotype to phenotype.

**Figure 2:**
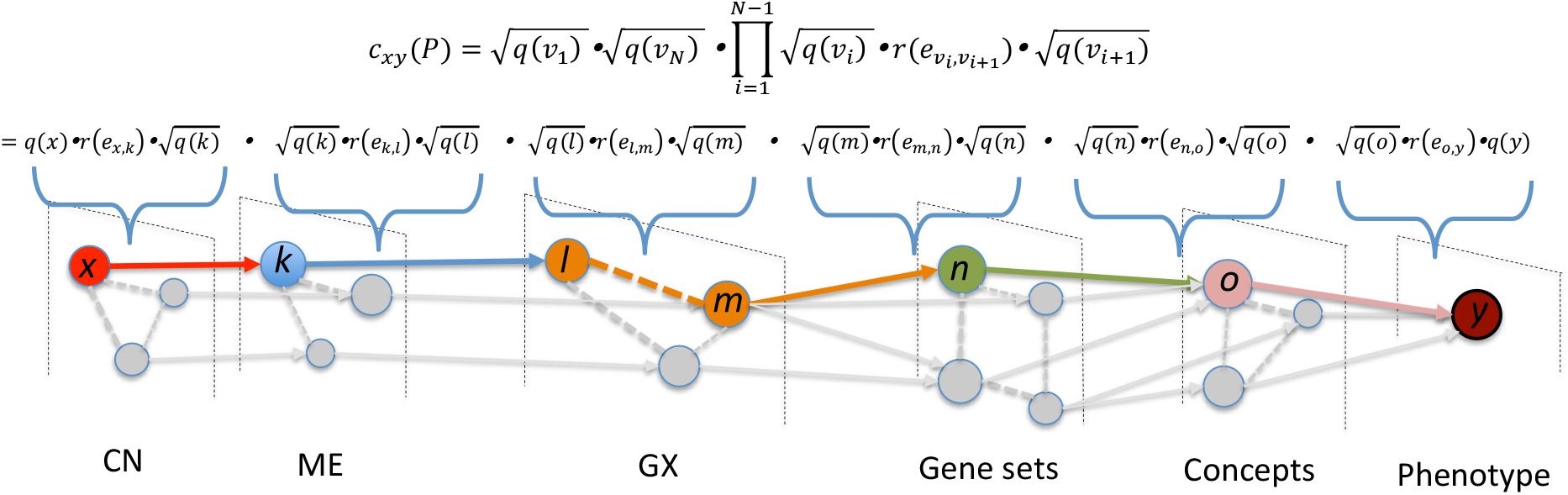
Maximum coefficient path analysis on multilayer network. *P* is an arbitrary path between *x* and *y* variables. The *path coefficient* is calculated by multiplying the correlation coefficients of all variables (nodes and edges) on that path.

## RESULTS

OmicsNet can be used for different kind of biological applications such as functional analysis, biomarker discovery or predictive analysis:

1. In *functional analysis*, the aim is to identify functional features, such as pathways or gene sets, that are most associated with the phenotype, while consistent with the genotype data and the multilayer structure. We can compute a *functional association* (FA) score (**Supplementary Methods Eq. (9)**) for each functional feature using the definition of the 3-point shortest path (**Supplementary Methods Eq. (7)**). Features with highest FA score will correspond to functional features that are mostly associated with phenotype and while being consistent with the genotype data.
2. In *biomarker discovery*, the aim is to identify genotype features that are most associated (“predictive”) with phenotype data, while consistent with the multilayer structure. Given a phenotype, and using the definition of the path coefficient, we can compute a *biomarker* score for each genotype feature (**Supplementary Methods Eq. (10)**). The biomarker score is simply the *path coefficient* from genotype feature to the phenotype.
3. In *predictive analysis*, the aim is to identify phenotype features that are most associated with some given genotype data, while consistent with the multilayer structure. Given a genotype, and using the definition of the path coefficient, we can compute a *prediction* score for each phenotype feature (**Supplementary Methods Eq. (11)**). The prediction score is simply the *path coefficient* from a specific genotype feature to the phenotype features. Predictive analysis of drug sensitivity is important for targeted therapy in personalized medicine.

The multilayer network approach elegantly unifies different types of analyses in a single framework. By conditioning the phenotype of interest in the multi-layer system, we can use OmicsNet for functional analysis, biomarker discovery or predictive analysis. In the following, we illustrate the application of OmicsNet in each type of analysis with examples in biology.

### Functional analysis: ABC-DLBCL versus GCB-DLBCL

Diffuse large B-cell lymphoma (DLBCL) is the most common subtype of non-Hodgkin lymphoma. There are two major biologically distinct molecular subtypes of DLBCL: germinal center B-cell (GCB) and activated B-cell (ABC) (Alizadeh et al. 2000). ABC-DLBCL is associated with substantially worse outcomes when treated with standard chemoimmunotherapy. ABC-DLBCL is defined by constitutive NF-kB pathway activation and BCR signalling pathways are oncogenically activated in this subtype: mutations in *MYD88, CARD11* and *CD79B* are found in ABC-DLBCL along with deletions and mutations of *TNFAIP3* (Pasqualucci and Dalla-Favera 2015). **Fig. 3** shows the multi-layer system comparing the two different subtypes ABC-DLBCL versus GCB-DLBCL. The entire system consists of seven layers, that includes four primary omics layers, two functional layers and a phenotype layer. Maximum coefficient paths were calculated for the ABC and GCB phenotypes separately. Systems analysis using OmicsNet showed that the ABC-DLBCL phenotype (in red) was characterized by concomitantly increased copy number, hypomethylation and overexpression of the genes *CD44, AHNAK, CYB5R2*, and *S100A6*. The system showed that the ABC-DLBCL phenotype is associated with activated IL-6 signalling, NF-kB pathway and the IL2/JAK/STAT program (in concordance with literature). On the other hand, the GCB-DLBCL phenotype (in green) is characterized by concomitantly increased copy number, hypomethylation and overexpression of the genes *MGMT, MME (CD10), CDKN2A, ALOX5*, and OAT, involved in E2F targets and cell cycle.

**Figure 3:**
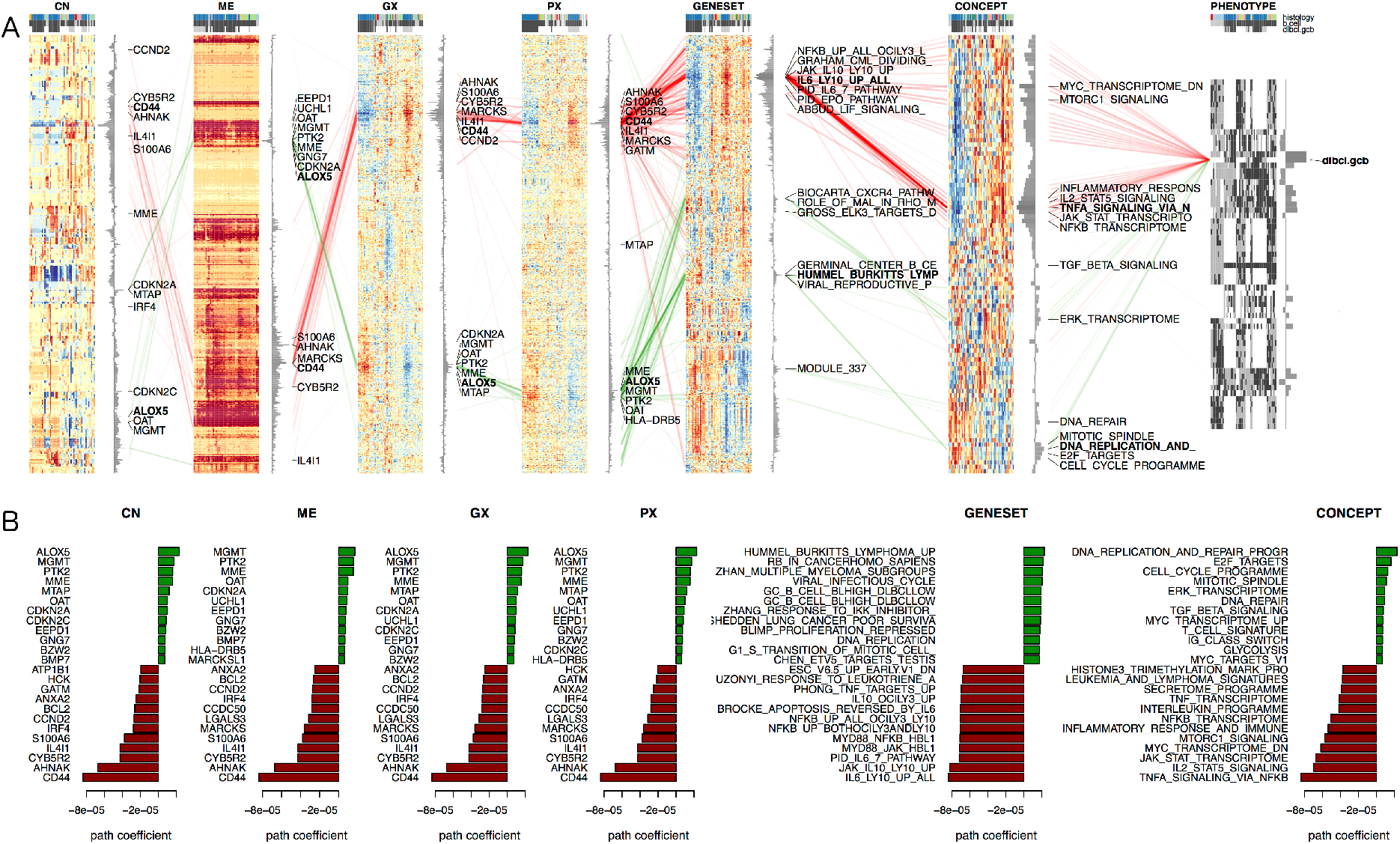
Functional analysis of ABC-DLBCL versus GCB-DLBCL. (A) Systems analysis using a 7-layer omics network. The maximum coefficient paths are highlighted in green for GCB-DLBCL, and in red for ABC-DLBCL. (B) Top scoring features with highest path coefficient for each layer. See main text for further details.

### Biomarker discovery: Sensitivity to Erlotinib in breast cancer

Erlotinib is tyrosine kinase inhibitor which acts on EGFR but has been shown to also be useful against HER2-overexpressing tumors (Schaefer et al. 2007). ERRB2 (HER2) is a member of the human epidermal growth factor receptor (HER/EGFR/ERBB) family. Amplification or over-expression of this oncogene has been shown to play an important role in the development and progression of certain aggressive types of breast cancer. Tamoxifen-resistant breast cancers often show increased expression of the epidermal growth factor receptor family members, ERBB1 and ERBB2. Baseline copy number, methylation and gene expression data of 35 breast cell lines (CCLE) were retrieved from the cBioportal. Single-sample gene set and concept features were calculated using rank correlation. Drug sensitivity values for 481 drugs were retrieved from the Cancer Therapeutics Response Portal (CTRP v2) database. We constructed a six-layer integrated network using the top 2000 most varying genes as it is shown in **Fig. 4**. Copy number (CN), methylation (ME) and gene expression (GX) layers are CIS connected. The system was solved for sensitivity to Erlotinib. The results confirm that response to Erlotinib is highly correlated with ERBB2 amplification (and overexpression). Sensitivity to the drug was associated with actived PI3K/MTOR/AKT transcriptome and cholesterol homeostasis. Genes *MIEN1, STARD3* and *MTAP* appear concomitantly amplified and overexpressed with ERBB2.

**Figure 4:**
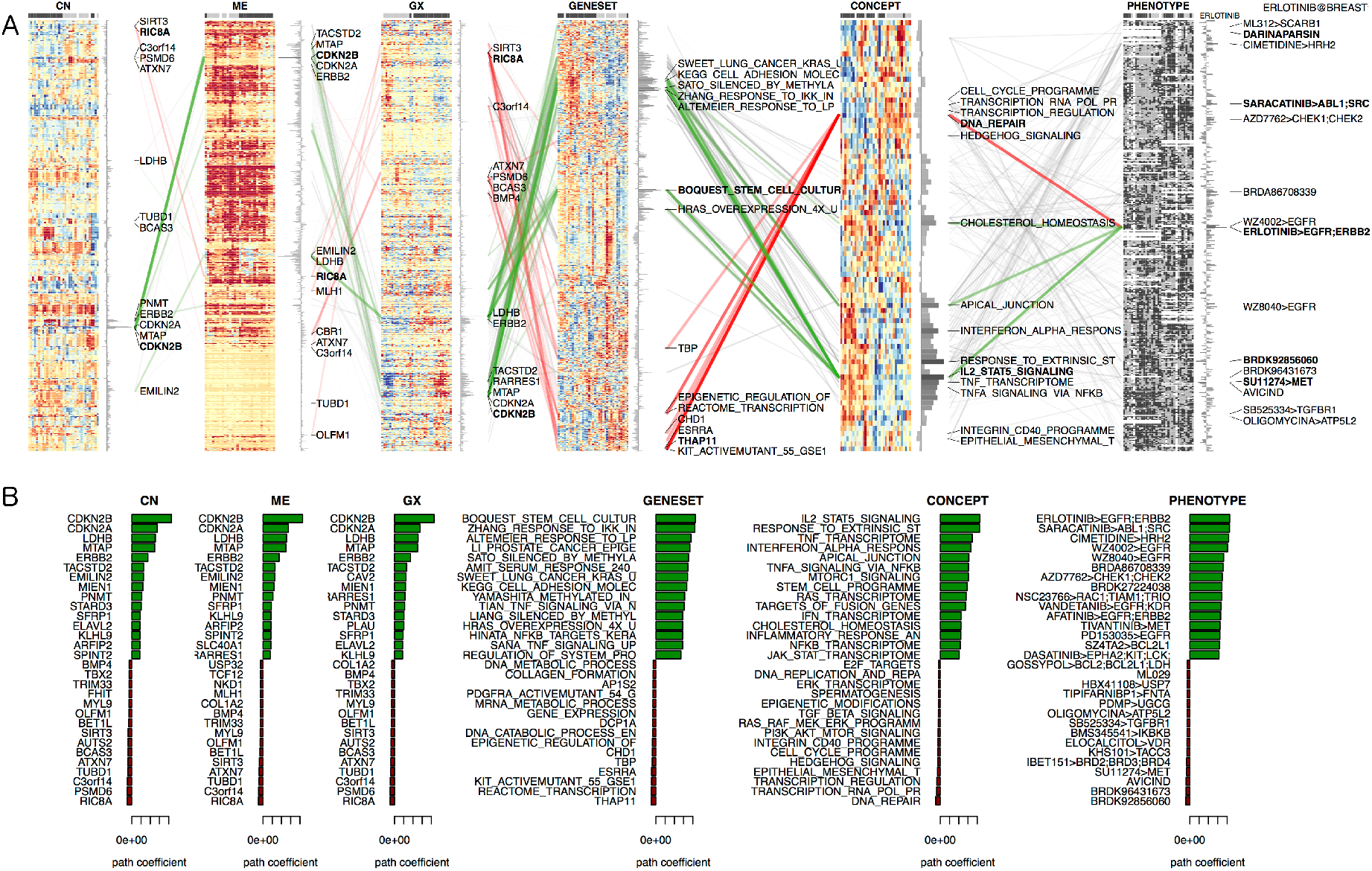
Biomarker analysis of Erlotinib sensitivity in breast cancer. (A) Systems analysis using a 6-layer omics network. The phenotype layer consists of drug sensitivity data was retrieved from CTRPv2. Maximum coefficient paths are highlighted in green for sensitivity, and in red for resistance to Erlotinib. (B) Top scoring features with highest path coefficient for each layer. See main text for further details.

### Predictive analysis: Targeted therapy for EGFR mutated lung cancer

EGFR has become an important therapeutic target for the treatment of these tumors. Inhibitors that target the kinase domain of EGFR have been developed and are clinically active. It is known that EGFR mutations activate the EGFR signaling pathway and promote cell-cycle (Zhang et al. 2010). Baseline mutation and gene expression data of 114 lung cancer cell lines (CCLE) were retrieved from cBioportal. Single-sample gene set and concept features were calculated using rank correlation. Drug sensitivity values for 481 drugs were retrieved from the Cancer Therapeutics Response Portal (CTRP v2) database. We constructed a five-layer integrated network shown in **Fig.5**. The mutation (MT) layer is fully connected to the gene expression (GX) layer. The system was solved for EGFR mutation as phenotype. Systems analysis showed that EGFR mutation in lung cancer was maximally correlated with the sensitivity to drugs such as saracatinib (targeting ABL1/SRC), gefitinib (targeting AKT1/EGFR), and WZ8040 (EGFR inhibitor). Mutant EGFR was associated with (as expected) increased EGFR transcriptome, higher glycolysis, and over-expression of genes including *TACSTD2, CD44* and *TGFBI*.

**Figure 5:**
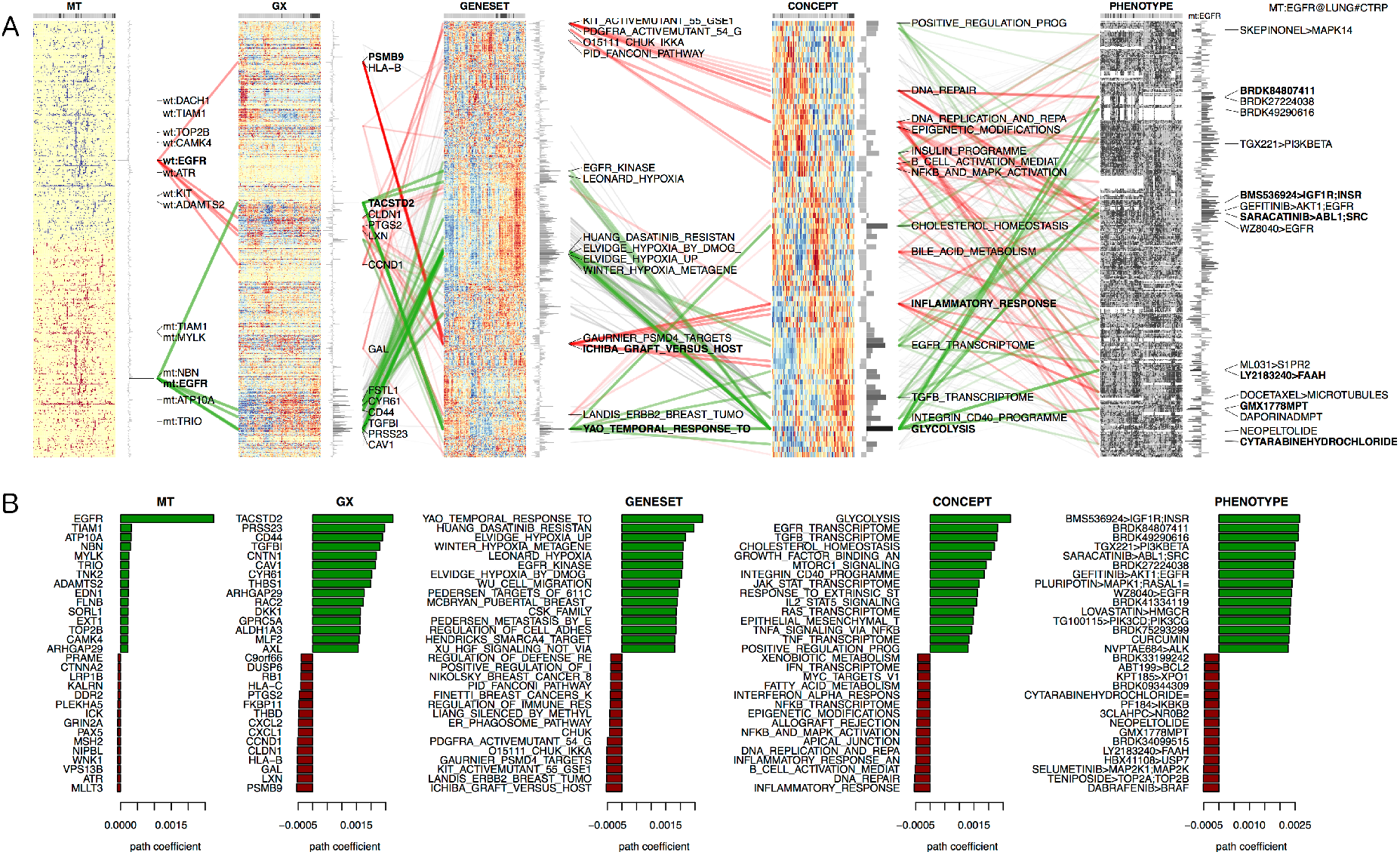
Predictive analysis of drug sensitivity for EGFR mutated lung cancer. (A) Systems analysis using a 5-layer omics network. Drug sensitivity was data retrieved from CTRPv2. Maximum coefficient paths are highlighted in green for mutated EGFR, and in red for wildtype EGFR. (B) Top scoring features with highest path coefficient for each layer. See main text for further details.

## DISCUSSION

OmicsNet is composed of input, intermediate and output layers, where input layers correspond to genotype features, intermediate layers represent geneset and biological concepts for functional interpretation, and output layer correspond to phenotypes. OmicsNet solves the integration problem by computing path coefficients between genotype and phenotype through the weighted multilayer network. OmicsNet naturally integrates and analyses multi-omics data based on ***path analysis*** in multi-layer networks. The variables on the maximum coefficient path have the highest joint correlation to genotype and phenotype, and biologically, may indicate the signaling cascade that is altered from perturbed features to a cellular response. The solution can be visualized as a multi-partite graph where the phenotype is connected to the most correlated omics and functional features. Quantifying the set of maximum coefficient paths is time consuming task and we transform this problem into finding the longest path problem, and later into the shortest path problem under our multilayer network topology, which computationally runs in linear time with respect to the number of variables.

The path coefficient can be regarded as a generalization of the scalar correlation coefficient to a path correlation, as a multiplication of correlation coefficients, quantifying the magnitude and significance of hypothesized causal connections between sets of variables along a path. While the scalar correlation of a feature indicates a local measure of association (to the phenotype), the path coefficient quantifies a global (integrated) fitness measure of multiple correlated features. A single feature may be highly correlated with the phenotype but may exhibit a low path coefficient if the overall connection to the phenotype is weak. In fact, the definition of the path coefficient as a multiplication means that all correlation values along the path must be large, and a single weak link immediately reduces the path coefficient.

The aim of systems biology is to develop a conceptual integration method to link the low level molecular perturbations to higher level phenotype. The structure of our multilayer network resembles the central dogma of biology of how genotype information is transcribed and translated into a phenotype. Accordingly, the input layers in our multilayer structure are composed of genotype information related to DNA or mRNA changes, and the output layer corresponds to phenotypes, whereas intermediate layers explain what type of biological pathways have been involved in the process. Intermediate layers facilitate functional interpretation of molecular mechanisms caused by changes in the genotype and leading to a particular phenotype. Currently, we use genesets and higher level concepts as intermediate layers. However, it is easy to extend it by adding already existing functional annotation databases as additional layers such as protein interaction networks and Gene Ontology (GO) graph. **Fig. 6** illustrates the use of the hierarchical Gene Ontology graph as a functional layer between the genomic and phenomic layers. Note the separation of input and output nodes in the GO graph, the low level GO terms serve as input and higher level GO terms act as output to the next layer.

**Figure 6:**
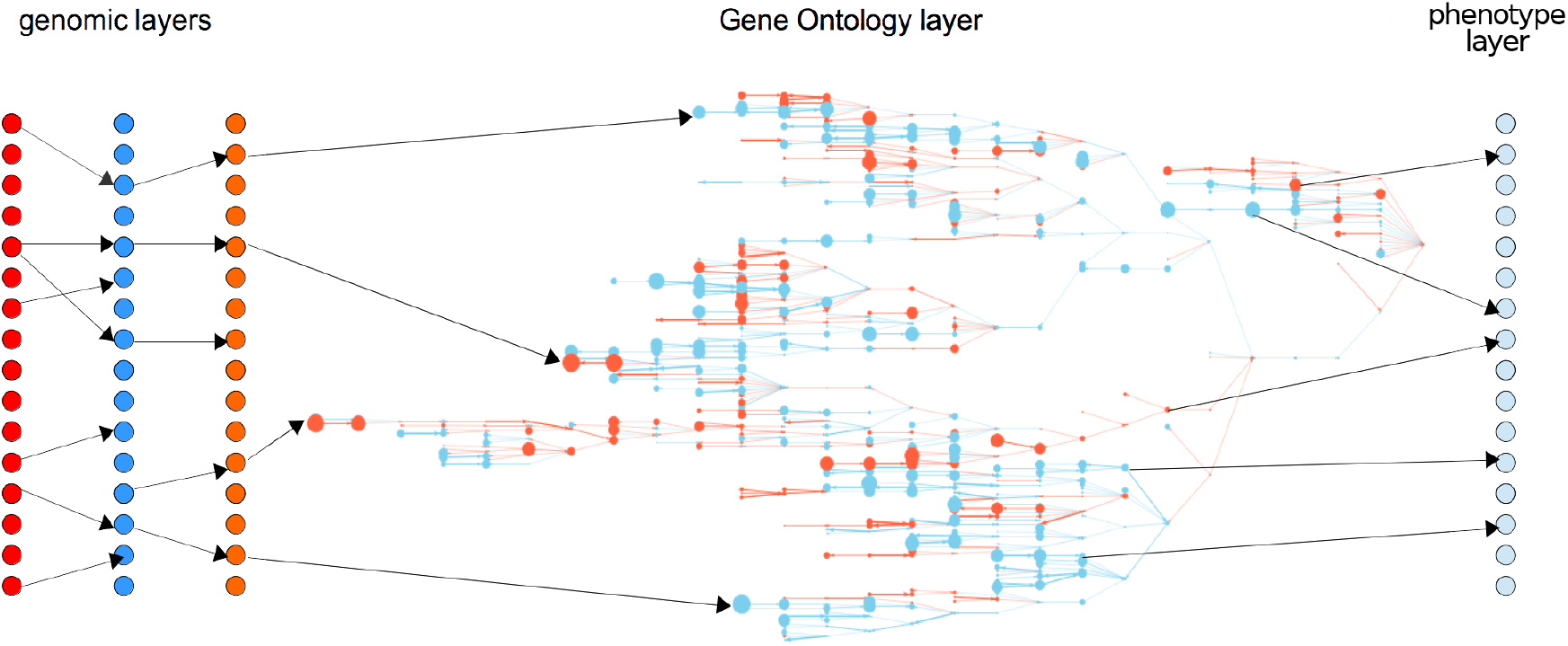
Illustration of a generalized multilayer network with three genomic layers, hierarchical Gene Ontology layer and final phenotype layer.

We have used the shortest path algorithm as our main solver for finding the maximum coefficient paths. Alternatively, the problem of finding the set of high coefficient paths in **Supplementary Methods Eq.(6)** can be formulated as a flow problem. Similar numerical solutions to the set of high coefficient paths satisfying **Supplementary Methods Eq. (6)** can be efficiently computed using the algorithms that search for an average or minimum cost flow. However, there are some other challenges associated with that. The edge weights and their upper limit flow capacities should be assigned before using the flow algorithms. One can use the inverse correlation between the variables as edge weights. However, it is not straightforward to adopt the notion of capacity in a biological context and assigning reasonable value to the capacities is not trivial. The solution structure of the minimum cost flow is highly dependent on the edge parameters.

There are some similarities and differences between OmicsNet and artificial neural networks (ANN). The multilayer structure of OmicsNet is similar to that of ANN, where both models have input, intermediate and output layers, that receive input, process it, and present the final result at the output layer. Notice the common ‘funnel’ structure, from input layers with high number of features to layers that have increasingly less features. While ANN might be considered as a black-box from the biologists point of view, OmicsNet is more transparent and easier to interpret because the intermediate features are known. In fact, the main purpose of OmicsNet is not prediction but *functional interpretation* of the data. Another difference is that ANN needs a training to be able to make classification or prediction analysis. In contrast, OmicsNet does not require any training to make functional interpretation, prediction and multi-phenotype testing. Currently, deep learning ANN models are not very appealing in omics data analysis due to the high number of variables versus small number of samples. Deep learning ANN would not be able to be trained with 10-100 number of samples currently typical in biological data sets.

## CONCLUSION

Integration of multiple omics data is becoming increasingly important to obtain a comprehensive view of dysregulated biological pathways in diseases and to better understand the underlying cellular mechanisms. The major challenge is how to properly integrate multi-omics data in a computational framework. In this paper, we introduced OmicsNet a novel method that uses a multilayer network for the integration and analysis of multi-omics data of heterogeneous types. In biological example applications, with typically five or more layers and hundreds of thousands of variables, our method successfully identifies genomic and functional modules that are related with the phenotype.

It is important to note that OmicsNet integrates multi-omics data in two aspects. Firstly, it integrates *multiple types* of omics data such as DNA copy number, mRNA, methylation, mutation, proteomics, and metabolomics. Additionally, it integrates *multiple scales* of data: from molecular data, to biological concepts, and finally to the macroscopic phenotype. The elegance of OmicsNet is that it integrates multi-omics and multi-scale features, all in a single multi-dimensional model.

The multilayer framework also unifies different types of analyses. We have shown how OmicsNet can be used for functional analysis, biomarker discovery and phenotype prediction of multi-omics data. Furthermore, formulating the omics data integration problem using multiplex graph structures allows the use of efficient graph optimization algorithms. Finally, by plotting the multi-layer graph, it is easy to visually analyze the causal effects between phenotype and genotype.

We believe our method will be significant contribution to available methods for multi-omics integration and be a starting point for a new class of integration algorithms based on multilayer graphs and path analysis.

## Acknowledgments

M.A. was supported by the Swiss National Science Foundation (205321-147138/1, www.snf.ch).

## Author contributions

I.K. and M.A. conceived of and designed the approach. I.K, A.A. and F.B. interpreted the biological findings. M.A., I.K and R.M. developed the mathematical aspects. I.K. and M.A. wrote the manuscript.

## Competing financial interests

A patent on the integration method as described in this paper has been filed as IPTO 102017000125434 on November 3^*rd*^ 2017.

## 1 SUPPLEMENTARY METHODS

### 1.1 The multilayer network as biological model

The use of multilayer networks in a biological setting has been previously in proposed (B. Wang et al. 2014) for clustering the patients using heterogeneous data. In this section, we define the multilayer framework used in our implementation for integrating omics data (see **Fig. 1**). Formally, we represent OmicsNet multilayer network as a tuple *G* = (*V, E, M*) where the set of vertices, edges and layers are represented by *V, E*, and *M* respectively. Each vertex *v_i_* ∈ *V* is assigned to exactly one layer *M_k_* ∈ *M* through some layer mapping function *f* : *v_i_* → *k*. A single edge connecting two vertices *v_i_* and *v_j_* will be denoted as *e_ij_*, while the set of edges connecting the vertices between the two layers *M_i_* and *M_j_*, is denoted as *E*_ij_. In most cases, each layer is comprised of a single data type such as gene expression, copy number, methylation, or gene set expression. Every node in the layers correspond to a feature, which is connected to other features in the same layer with an intralayer edges and linked to features in other layers with an interlayer edges.

The order of the layers in the network crucial for correct modelling and is based on biological knowledge. Edges are generally undirected in correlation-based multilayer networks, but the direction of the edges between layers is generally imposed by the assumed multilayer structure. In biological applications, we assume that information in the multilayer network propagates from the “genotype” to “phenotype” layers. It is known that the copy number alteration and methylation status control the gene expression levels. Different biological scenarios can be employed to formulate the relation between the copy number and methylation layers. We modeled the relationship between the copy number and methylation using *or* operator. Another option could be to merge these two layers, or add an artificial layer in front of them to coordinate the causality relationship. We also add geneset and biological concepts layers to identify higher level functional features and pathways that associated with the variability in cellular responses. Quantitative values for geneset and concepts are computationally inferred for each single sample from the primary measured omics data types. The biological concepts are higher-level terms for functional analysis and are composed of gene sets, further reducing the feature space and thus simplifying the functional interpretation.

The proposed multilayer network structure is general and additional omics data types can be easily incorporated by constructing additional layers. For instance, the proteomic, phosphoproteomic, and metabolomic data can be added between the gene expression and gene set layers since they harbor post-transcriptional and post-translational biological information closer to the phenotype.

### 1.2 Path coefficient analysis in multilayer networks

Multilayer networks represent the “genotype-to-phenotype” relation in biological systems, where the variables in input layers (‘genotype’) have effect on output layer (‘phenotype’) through intermediate layers. The aim is to quantify the relationship between all genotype variables, the variables in the intermediate layers and ultimately the phenotype variables in the output layer. In order to validly calculate this relationship, Wright (1934) proposed a simple set of path tracing rules for calculating the coefficient correlation between two variables, which is equivalent to the total contribution of all the pathways that connect two variables. So called the “product rule” states that the net causal impact of a path connecting two variables *x* and *y* is the product of the correlation of all variables in between. This quantitative measurement is called the *path coefficient*, and it is symbolized by *c_x_y* for the causal impact of *x* on *y*.

Now, imagine *x* and *y* represent some variables in genotype and phenotype layers of **Fig. 1**, respectively, and the ordered layers of network are denoted by *M* = (*M*_1_, *M*_2_,…, *M_n_*). Let *P* = (*v*_1_, *v*_2_, *v*_3_,…, *v_t_*) be an arbitrary path sequence of a number of *t* vertices such that *v_i_* is linked to *v*_*i*+1_ through an edge *e*_*i,i*+1_ for 1 ≤ *i* ≤ *t*, and *x* = *v*_1_ and *y* = *v_t_*. Each vertex *v_i_* is assigned to a layer *M_k_i__* through layer mapping function *f* : *v_i_* → *k_i_*, such that *k_i_* <= *k*_*i*+1_. The latter states that two consecutive vertices of the path *P* can be within the same layer, or can traverse to the next layer but cannot travel backwards to any previous layers. Each edge carries a weight 0 < *r*(*e*_*i,i*+1_) ≤ 1 that is equal to the correlation coefficient between its incident variables. We generally only consider positively correlated edge weights since it better interprets the relationship between features and phenotype. In certain cases, one may consider the absolute value of the negative part of the correlation value in the computations while modeling negatively correlated biological information flow. Instead, the absolute correlation value may be used as edge weights when the sign of the biological information flow is deemed not to be relevant.

The prize of the vertex *v_i_* is denoted by *q*(*v_i_*), and generally quantifies some perturbation of that variable with respect to the phenotype. For instance, the prize of a vertex can be a copy number difference, expression fold change, number of mutations, or any parameter that is correlated with the phenotype. The prizes are required to be strictly between 0 < *q*(*v_i_*) ≤ 1 so we generally take the absolute value of the parameter or perform the path analysis for positive and negative node prizes separately. We found that the latter is preferred, in order to find coherent features in relevant biological functions. Then, the *path coefficient* between *x* and *y* is computed as follows:

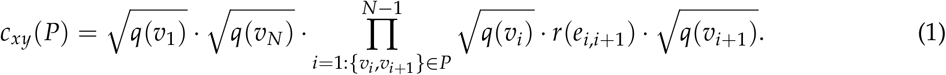

A simple example for the computation of the path coefficient is illustrated in **Fig. 2**. After properly defining the *path coefficient* of a single path, we analyze all possible paths between genotype and phenotype layers in the multilayer network, and identify the paths that have the maximum coefficient values. The variables on these paths have the highest joint correlation to both the genotype and phenotype simultaneously, and biologically speaking, may indicate the solution that best integrates multiple variables associated with the phenotype. Let *P*_all_ be all the possible paths from the genotype variable to phenotype variable, then the target is to detect the collection of paths *S*(*P*_max_) such that:

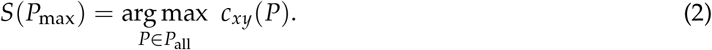

Since the number of paths with maximum coefficient could be very limited, the variables on those paths may not be sufficient to explain the difference in cellular response. Alternatively, one may want to examine the “neighbourhood” of the maximum coefficient solution to evaluate robustness of the solution, or to explore alternative near optimal paths that may improve the interpretation. To this end, we search for the set of paths *S*(*P_set_*) with coefficient value higher than a threshold *T* as follows:

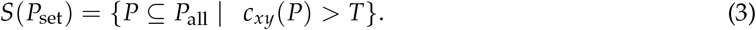

### 1.3 Computing the path coefficients using the shortest path

There are finitely many paths between the variables of input and output layers in multilayer network, and we need an efficient method to enumerate all the paths and compute their coefficient values. While multiple paths are possible, the maximum coefficient path gives the best quantitative measure of how likely the causal relation is. We can rewrite **Eq. (2)** for detecting the maximum coefficient path using the logarithmic expression:

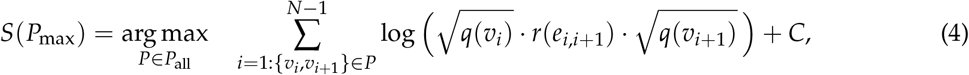

where the constant term corresponds to 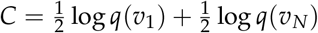. Now, it is obvious to see that **Eq. (4)** is very similar to the definition of the *longest path* from the vertex of the input layer to the vertex of the phenotype considering the loop constraints of the paths. Since the terms inside the logarithm are always smaller or equal to one, their logarithmic value must be negative and therefore by negating the term inside the summation, we can use the *shortest path* (Dijkstra 1959) to effectively find the maximum coefficient path:

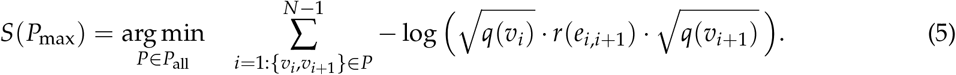

In order to compute the coefficients of all the possible paths, one can solve *all-pairs shortest paths* from the vertices of input layers to the phenotype nodes. Similarly, **Eq. (3)** can be rewritten to find the set of paths with the coefficient value higher than the threshold using the *shortest path* analogue:

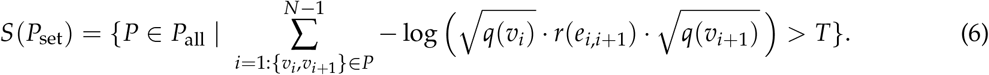

### 1.4 Path coefficients from three-point shortest path solution

We generalize shortest path problem to find the shortest path between three points. Let *P_xyz_* = (*v_x_*,…, *v_y_*,…, *v_z_*) be a viable path in the multilayer network that that starts from vertex *x*, traverses vertex *y* and reaches vertex *z*. We define the “3-point path coefficient” *c_xyz_* as a product of path coefficients:

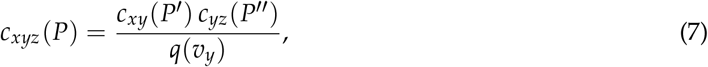

where *P*′ = (*v_x_*,…, *v_y_*) and *P*″ = (*v_y_*,…, *v_z_*) are the partial sections of the entire path *P* to and from vertex *y*, respectively. Then using **Eq. (2)** we can retrieve the set of “maximum 3-point coefficient” paths *P_xyz_* by solving the following maximization problem:

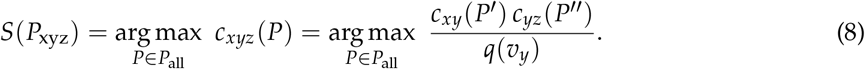

The equation above can be solved by finding the arguments of the maximima of the two terms individually. In other words, to find the 3-point shortest path *P_xyz_* we can solve for the shortest paths *P_xy_* and *P_yz_* separately, and concatenate their solution. The 3-point path coefficient is then simply the product of path coefficients as given in **Eq. (7)**.

### 1.5 Characteristics and generalizations of the method

- A numerical solution of **Eq. (5)** can be efficiently computed by Dijkstra’s algorithm solving for single-source shortest path (SSSP). A similar numerical solution for the set of shortest paths that satisfy **Eq. (6)** might be efficiently obtained using algorithms that compute the average or minimum cost flow. Given points *x* and *y*, the 3-point shortest paths in **Eq. (7)** for all vertices *y* can be computed efficiently by solving the SSSP twice from *x* and *z*.
- A shortest path (SP) solution between nodes in the input and output layer traverses each layer of the multilayer network at least once. In layers that have intra-layer connections, it is possible that the SP contains multiple traversals *within* a single layer.
- Input layers generally correspond to a measured (omics) data types. Intermediate layers generally correspond to other measured data types, or are computationally derived from measured data types such as the gene set and concept layers. The final output layer (or ‘phenotype layer’) generally corresponds to sample phenotypic information such as histology or cell type, or clinical information in the case of a patient, such as cancer subtype, drug response, or survival.
- Layers can have both intralayers connections (connecting nodes within the layer) and interlayers connections (connecting to different layers). For a particular layer one may disallow intralayer connections to model *independency* between features. One may restrict the number of intralayer connections to improve computational and memory efficiency. Between the omics layers, interlayer connections may be restricted to nodes that belong to the same gene/protein to model CIS-regulation between data types.
- Note that the phenotype layer can contain multiple phenotypes at once. Thus the multilayer model can be used for modelling of multiple phenotypes. Intralayer connections in the phenotype layer may model *multicollinearity* in the phenotype information. Note that the correlation values *r*(*e*) on the edges are independent of the selected phenotype but the vertex prizes *q*(*v*) not.
- The number of layers is not limited and depends on the number of different data types and functional layers in the model. Currently, the configuration (order) of the layers have to be decided. Perhaps it will be possible to infer the optimal configuration of the layers from the data. Recurrent layer connections (or loops) are not allowed in our formulation because the shortest path cannot handle that. Perhaps by modeling the signals dynamically, or using Monte Carlo type of solvers, one can model recurrent connections.
- A layer in the multilayer network may specify distinct input and output nodes such that the connections to the previous layer and to the next layer are forced to be different nodes. **Figure 6 illustrates the hierarchical Gene Ontology graph as a generalized layer with different input and output nodes. By specifying non-overlapping input and output nodes, the shortest path solution is forced to traverse within the layer. For the genomic layers, one possibility could be to specify signalling genes or kinases as input, and transcription factors as output of the layer.
- Dimensions of the layers are proportional to the number of data features and depend on the measurement technology used. Omics data layers comprise several 10’000 nodes corresponding to genes or proteins. Gene set layers typically contain 1000 to several 10’000 of nodes. Concept and phenotype layers typically contain hundreds of variables.
- External interactome databases, such as protein-protein interaction networks, gene ontology, gene signalling pathways, or gene regulation networks, may be used to construct or weight the intralayer connectivities of the appropriate data layer. Fig. 6 illustrates using the gene ontology graph as a functional layer between the genomic and phenomic layers.
- It is also possible to have a multiple layers corresponding to the same data type, for instance, if the data is obtained as time series at different points. The time series data can be fused into one layer, or can be used as it is.
- In certain applications, it is convenient to introduce “source” and/or “sink” layers each with a single vertex, placed before the input layers and behind the output (i.e. phenotype) layers.

### 1.6 Functional analysis using the multilayer system

Given a multilayer graph *G* = (*V, E, M*), we differentiate the layers into the set of input layers *M_x_*, intermediate layers *M_y_*, and output layers *M_z_*. We refer to input variables in *M_x_* as *X*, features in the intermediate layers *M_y_* as *ϒ*, and output phenotype variables in *M_z_* as *Z*. Note that *ϒ* may include both omics and functional features, depending on the multilayer model. Furthermore, we partition the set of corresponding vertices accordingly into *V* = (*V_x_, V_y_, V_z_*) where *V_x_, V_y_* and *V_z_* denote the set of vertices corresponding to the input features *X*, intermediate features *ϒ*, and output phenotype features *Z*, respectively.

In functional analysis, the aim is to identify those functional features *ϒ* that are most associated phenotype data *Z*, while consistent with the genotype data *X* and the multilayer structure *G* = (*V, E, M*). A straightforward method is to computing the shortest path (SP) solution *S*(*P*_max_) as in **Eq. (5)**, or the set of SP solutions *S*(*P_s_*) as in **Eq. (6)**, and take the intersection those features with features that are in *V_y_*. The drawback of this method is that we do not have a score for features *v* ∈ *V_y_* that are outside of the SP solution set.

Alternatively, given a phenotype *z* ∈ *V_z_*, using the definition of the 3-point shortest path in **Eq. (7)**, and by introducing source node *s* connected to all vertices in the input layers, we can compute a *functional association* (FA) score for each functional feature *y* ∈ *V_y_* as

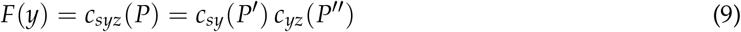

where *P*′ and *P*″ are the partial paths as in **Eq. (7)**. That is, the functional association score *F*(*y*) for feature *y* is simply the 3-point shortest path from a virtual “source” vertex *s*, through *y*, to phenotype *z*. Features with highest FA score will correspond to functional features that are most associated with phenotype *z* and while consistent with the data *X*. Note the addition of an extra layer to include the *source* node *s*.

### 1.7 Biomarker analysis using the multilayer system

In biomarker analysis, the aim is to identify genotype features in *X* that are most associated with phenotype data *Z*, while consistent with the multilayer structure *G* = (*V, E, M*). Given a phenotype *z* ∈ *V_z_*, and using the definition of the path coefficient in **Eq. (1)**, we can compute a *biomarker* score for each *x* ∈ *V_x_* as

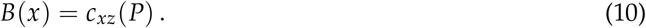

That is, the biomarker score *B*(*x*) for genotype feature *x* is simply the *path coefficient* from *x* to the phenotype variable *z*. Note that the exact shortest path between *x* and *z* is constrained by the structure of the multilayer network, and generally must traverse any other omics layers and higher order functional layers. Thus such a score integrates the association of all intermediate variables.

### 1.8 Predictive analysis using the multilayer system

In predictive analysis, the aim is to identify phenotype features in *Z* that are most associated with genotype data *X*, while consistent with the multilayer structure *G* = (*V, E, M*). Given a genotype *x* ∈ *V_x_*, and using the definition of the path coefficient in **Eq. (1)**, we can compute a *prediction* score for each *z* ∈ *V_z_* as

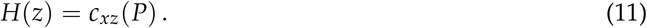

That is, the predictive score *H*(*z*) for phenotype feature *z* is simply the *path coefficient* from genotype *x* to the phenotype variable *z*.

### 1.9 Meta-analysis of multiple phenotypes

Let *G* = (*V, E, M*) be a multilayer network, and let *K* be multiple phenotypes encoded by *Z* = (*z*_1_, *z*_2_,…,*z_K_*). For each phenotype *z_i_*, we compute the *biomarker score* for all features *V_x_* in the input layer, and *functional association* (FA) score for all features *V_y_* in the intermediate layers, according to **Eq. (10)** and **Eq. (9)**, respectively, such that

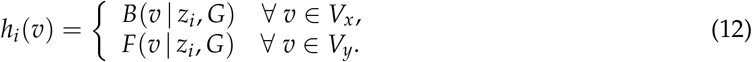

Then, the matrix *H* = (*h*_1_, *h*_2_,…, *h_K_*) contains biomarker scores or FA scores for all features *v* ∈ (*V_x_, V_y_*), for each phenotype in *Z*. One can compare pairs (or groups) of *h_i_* to detect commonalities or differences in phenotype associated features depending on the phenotype.

### 1.10 Statistical significance

P-values for assessing the statistical significance of the path coefficients, can be computed numerically by permutation of the labels and randomization of the node and edge values of the network.

## 2 IMPLEMENTATION NOTES

### Calculation of geneset summaries

The functional layers need either gene set or concept summaries to be calculated for each sample. Single-sample gene set methods such as GSVA or ssGSEA can be used. Especially for large number of gene sets, we have implemented a moderated rank correlation measure that is very efficient to calculate and retains good agreement with GSVA and ssGSEA. Gene sets were retrieved from MSigDB and Harmonizome. Concepts were calculated from the “founder” sets of the hallmark gene sets of MSigDB, plus additional biological concepts manually curated by ourselves.

### Sign of edges

For reasons of interpretability of the path solution, we generally are only interested in edges with *positive* correlation. With exception of the *methylation* layer that is forced to be negatively correlated with the *transcriptome* layer, and with the exception of the *mutation* layer allowing both positive and negative edge value (i.e. correlation) to the *transcriptome* or *proteome* layer.

### Memory size

Ideally, the multilayer graph would be fully connected, i.e. all features within layers and between layers would be connected with each other. However, this would quickly lead to memory allocation problems because of the huge number of edges. In R, a dense correlation matrix of either within a single layer or between two layers with 20’000 features takes approximately 3.2Gb of memory space (8 bytes per double precision float). The corresponding igraph object took 2.34Gb of memory space. A fully-connected multilayer graph with several layers would quickly exhaust computer memory of a typical PC. To reduce the overall required memory, but also to improve the computational efficiency, we enforce a number of contraints while building the multilayer graph:

1. The number of features in the layers are limited by taking the top *N* most varying features (ranked by decreasing SD). We estimate that *N* = 4000 suffices for most cases. For quick evaluations we use *N* = 1000. Data sets with many tissue types may need more features, for example *N* = 8000, to capture the heterogeneity.
2. Intra- and interlayer connections are limited to the top *N* edges with highest correlation. By default we set a maximum of *N* = 20 for intra-connections, and a maximum of *N* = 100 for interlayer connections. The maximum number of allowed intra-connections is lower because we expect the shortest path solutions not to traverse many steps within the layers.
3. Interlayer connections between genomics layers are CIS constrained, meaning inter-layer connections are allowed only between the same gene. This considerably reduces number of edges. For the *mutation* layer, we do not apply a CIS constraint because, in general, a mutation may not have an effect to the mRNA transcriptome expression, unless it is a truncating mutation.
4. Copy number, methylation and mutation layers are assumed to have *independent features*. As such, these layers do not allow intra-layer connections. This further reduces the number of edges.

### Graph implementation

The multilayer graph model is implemented using the igraph R package. As explained above, we avoid creating a fully-connected multilayer graph. As such, instead of creating the graph from a dense correlation matrix, the graph is instantiated using a custom edge list that respects all constraints in order to control the sparsity of the graph at construction.

### Shortest path calculation

Shortest paths (SP) are computed using the shortest_paths function of the igraph R package based on the the Dijkstra algorithm. We are currently also testing a GPU-based SP solver to accelerate the computations especially when computing permutation p-values. For each phenotype we compute two shortest paths corresponding to the ‘positive’ and ‘negative’ phenotype seperately. The ‘positve’ solution comprises of the shortest path that traverses only node with positive node prizes, i.e. features that are striclty positive correlated with the phenotype. Conversely, the ‘negative’ solution comprises of the shortest path that traverses only node with negative node prizes, i.e. features that are negatively correlated with the phenotype.

